# Modeling electron interference at the neuronal membrane yields a holographic projection of representative information content

**DOI:** 10.1101/2022.12.03.518989

**Authors:** Elizabeth A Stoll

**Affiliations:** Western Institute for Advanced Study, Denver, USA

**Author notes:** https://westerninstitute.org.

## Abstract

It has historically proven difficult to explain the relationship between neural activity and representative information content. A new approach focuses on the unique properties of cortical neurons, which allow both upstream signals and random electrical noise to affect the likelihood of reaching action potential threshold. Here, each electron is modeled as an electromagnetic point source, interacting in a probabilistic manner with each neuronal membrane. The electron is described as some set of probability amplitudes, distributed across five orthogonal axes: *x, y, z, energy state*, and *time*. The membrane potential of each neuron is defined by the probabilistic spatial position and atomic orbital of each local electron, after some time evolution. The mixed sum of all probabilistic component pure states is the physical quantity of information held by the neural network, given by a complex-valued wavefunction. If the probabilistic trajectory of each electron over time *t* affects the voltage state of multiple computational units, then the system state must be computed as a whole, with the state of each neuron being resolved as every component pure state is resolved. This computational process yields a defined system state at a defined location in time, which immediately becomes the past as a new probability density forms. If the membrane surface of each computational unit is also a charge-detecting polymer substrate that meets the criteria of a holographic recording surface, then this encoding process will generate a holographic projection of representative information content. The constructive and destructive interference of high-dimensional probability amplitudes yields a non-deterministic computational outcome for each neuron. That now-defined system state is paired with a multi-sensory percept, which is exclusively accessed by the encoding structure, with content limited by the range and sensitivity of the sensory apparatus. This model usefully offers a plausible explanation for both perceptual content and non-deterministic computational outcomes emerging from cortical neural network activity.

## I. INTRODUCTION

Over the past 100 years, the field of neuroscience has demonstrated that perceptual experience is correlated with neural activity in the cerebral cortex. This relationship has been established through both imaging and electrophysiological studies [1–4]. Data from the local environment are collected by the sensory apparatus and processed in the central nervous system [5, 6]. The resulting neural firing patterns across the cerebral cortex are correlated with qualitative perceptual awareness, the formation of memories, and the initiation of contextually-appropriate behavior [7–10]. Yet while neuroscience has made enormous strides in understanding the neural correlates underpinning perception, cognition, and decisionmaking, an explanatory gap persists. To date, there remains no plausible mechanistic explanation for how a bound perceptual experience might arise from neural network activity, nor how mental representation might participate in information processing.

To address these issues, it may prove useful to model the unique characteristics of cortical neurons. To engage in signal propagation, neurons set up electrochemical potentials, with a high concentration of sodium ions outside the cell and a negative voltage across the cellular membrane. The binding of excitatory neurotransmitters at chemical synapses permits the inward flow of positively-charged sodium ions, raising the membrane potential and triggering the opening of voltage-gated ion channels [11]. If a neuron reaches a certain voltage threshold prior to rectification of the membrane potential, the cell fires an action potential, releasing neurotransmitter to post-synaptic neurons. In invertebrate neural circuits and spinal reflex circuits, signaling outcomes for individual neurons are deterministic, and can be easily predicted by summing upstream inputs [12, 13]. Meanwhile, signaling outcomes for cortical neurons are highly unpredictable [14, 15]. Cortical neurons allow stochastic ion leak and spontaneous subthreshold fluctuations in membrane potential to affect the likelihood of firing an action potential [16, 17]. Uniquely, cortical neurons do not encode information in the spikes themselves, but rather in the probability of a spike occurring.

It has been proposed that ‘concepts’ are represented by unique patterns of activity in spiking neural networks [18], but no pattern in spike timing, spike rate, or phase coding has ever produced a plausible mechanistic explanation for the generation of representative information content [19]. A leaky integrate-and-fire (LIF) model is also commonly used to approximate cortical neuron signaling outcomes [20], but this approach does not effectively model the random firing patterns of cortical neurons, which act more as coincidence detectors than temporal integrators [21]. During a cortical up-state, when many neurons across the network are residing right near action potential threshold, the contribution of random electrical noise is sufficient for individual neurons to switch from an off-state to an on-state, leading to inherently probabilistic signaling outcomes [16, 17]. Since random electrical noise reduces both accuracy and energetic efficiency in classical circuits, cortical neurons are either less accurate and less efficient binary computing units than spinal neurons, or they are allowing random electrical noise to drive a non-deterministic computation.

A new approach focuses on the latter hypothesis, by modeling each electron within the system as a single electromagnetic point source, interacting probabilistically with every region of neural membrane. The neuron combines upstream signals with this random electrical noise to gate a signaling outcome. Here, the outer membrane of each computational unit acts as a charge-detecting polymer recording surface, both physically encoding information and generating a holographic projection of the encoded information content. This approach takes into account both random and non-random events contributing to cortical neuron signaling outcomes, and in doing so, it provides a mechanistic explanation for how cortical neurons generate both perceivable information content and non-deterministic computational outcomes.

## II. METHODS

### A. Modeling the cortical neuron as a two-state quantum system

During up-state, cortical neurons linger at their action potential threshold, allowing both upstream signals and random electrical noise to prompt a signaling outcome. So, while a neuron is classically interpreted as a binary logic gate in an ‘on’ or ‘off’ state, coded as 1 or 0, it could also be described as having some *probability* of converting to an ‘on’ state or remaining in an ‘off’ state. In this approach, a cortical neuron integrates upstream signals with random electrical noise, defining its voltage state as a function of time, as the system is perturbed. The neuron starts in off-state *ϕ*, not firing an action potential, and over time t, it reaches another state *χ*. And so, over some period of time, from *t*_0_ to *t*, the state of the neuron evolves from *ϕ* to *χ*. The timepath taken from one state to another is given by:

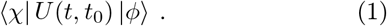

The probability of a state change can be represented in some basis:

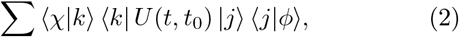

Such that *U* is completely described by base states *k* and *j*:

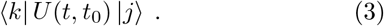

The time interval can be understood as being *t* = *t*_0_ + Δ*t*, so identifying the state of the neuron *χ* at time *t* can be understood as taking a path from one state to another:

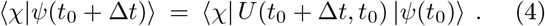

If Δ*t* = 0, there can be no state change. In this case:

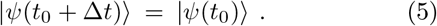

In any other case, the state of the neuron at time *t* is given by the orthonormal base states *k* and *j*, with probability amplitudes:

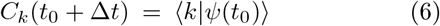

And:

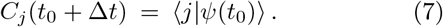

The state vector *ψ* at time *t* is a superposition of the two orthonormal base states *k* and *j*, with the sum of the squared moduli of all probability amplitudes being equal to 1:

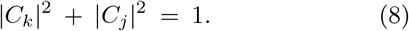

The neuronal state |*ψ*〉 at time *t* can therefore be described as a normed state vector *ψ*, in a superposition of two orthonormal base states *k* and *j*, with probability amplitudes *C_k_* and *C_j_*. Since the neuron starts the time evolution in state |*ψ*(*t*_0_)〉 = *j*, its probable state at time *t* is given by:

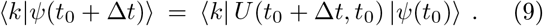

This equation can also be written in expanded form as the sum of all transition probabilities:

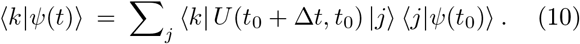

For the state vector *ψ*(*t*), the probability of a state change at time *t* is described by the U-matrix, *U_kj_*(*t*):

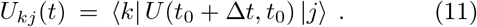

And so, all probability amplitudes are dependent on the amount of time that has passed, Δ*t*:

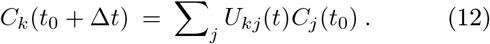

If Δ*t* = 0, there can be no state change and *k* = *j*. If Δ*t* > 0, there is some probability of a state change, where *k* ≠ *j*. As such, the two-state quantum system is described by the Kronecker delta *δ_kj_*:

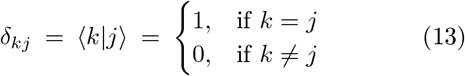

Here, the neuronal state |*ψ_n_*〉 evolves over time, with a binary signaling outcome at time *t* being a function of all local ion states. The state of each ion in the system |*ψ_i_*) also evolves with time, with its location at time t defined in relation to each local neuron, either inside that neuron or outside it. Any change in the location of a particular sodium ion is a function of the electrochemical potentials of all nearby neurons, which affect the activation state of all local ion channels, and the position, momentum, and energy state of every component electron. If the fundamentally uncertain state of an electron affects the state of the entire ion, in the presence of a dynamic electrical field, the present state of that ion is probabilistic, and the resultant voltage state of each neuron is also probabilistic. This uncertainty is expected to affect the membrane potential of cortical neurons in up-state, with quantum events actually contributing to the probability of a signaling outcome.

### B. Modeling the interactions between electrons and the neuronal membrane

Every electron in the system has some possible energy orbital *ω*, and some position in the *x,y,z* plane, both of which are fundamentally uncertain at time *t*. Any energy above ground state may be dissipated toward the production of *information*: a distribution of complex-valued probability amplitudes describing the possible states of each electron. In modeling the electron, the state vector *ψ* can refer to its energy orbital, which can be any one of several orthonormal pure states |*ψ_e_*〉.

Since the neuron’s membrane potential is dependent on the location of each electron in the system, and the location of each electron is uncertain, the neuronal membrane potential is also uncertain. A cortical neuron can therefore be considered a two-state quantum system, with some probability of undergoing a state change over time *t*. The cortical neuron starts the time evolution below the threshold for firing an action potential, in state *ϕ_n_*, but has some probability of reaching that threshold over time *t*, as it evolves into state *χ_n_*. The state of the neuron at time *t*, either having reached the threshold for firing an action potential or not, is represented by the state vector *ψ_n_*, and exists in a superposition of two orthonormal base states with probability amplitudes *C_k_* and *C_j_*.

This model abandons the classical tradition of considering a cortical neuron as a binary computing unit, always in an on-state or an off-state, firing an action potential or not, at any given moment. Instead, it considers the cortical neuron as a two-state quantum system, described by the Kronecker delta. Here, the computational unit calculates the *probability* of firing an action potential, as a function of all probabilistic component states. Since a cortical neuron allows random electrical noise to gate a signaling outcome [16, 17], and each electron may contribute to the voltage state of multiple neurons, the state of each neuron must be computed simultaneously, as every component pure state is computed. In this approach, the information that is physically encoded by each cortical neuron is a probability distribution - the mixed sum of all component pure states, or the von Neumann entropy of the system.

### C. The two assumptions underlying this theoretical approach

This model makes two theoretical assumptions: 1) cortical neurons permit random electrical noise to affect the likelihood of firing an action potential, and 2) the charge-detecting polymer membrane of each computational unit meets the criteria of a holographic recording surface. If this model is true, then 1) both of these assumptions should be empirically verifiable, and 2) these two key features should be both necessary and sufficient to produce perceivable information content. The justification for making these two assumptions is given below.

### D. Justification for modeling the membrane potential as a mixed sum of component pure states

One assumption is that cortical neurons allow random noise to affect signaling outcomes, particularly during a cortical up-state, when many cells across the network are poised at action potential threshold. The justification for this assumption is provided in Figure 1.

**Figure 1.**
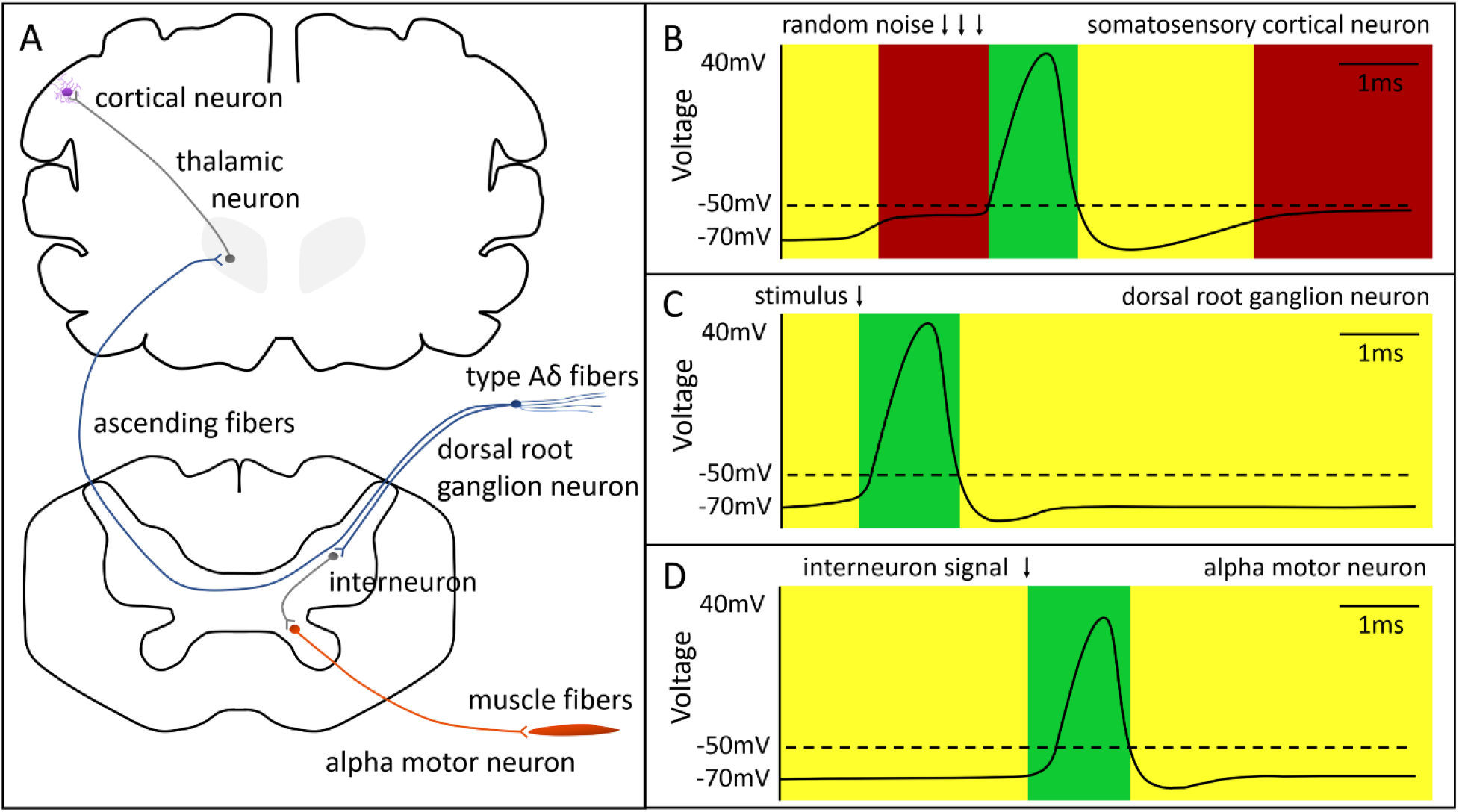
The unique physiological feature of cortical neurons: engaging in probabilistic coding. The proposed theoretical model differentiates between spinal reflexes, which are *deterministic* and *not correlated with a cohesive stream of perceptual experience*, and cortical circuits, which are *sensitive to random electrical noise* and *correlated with representative information content when processing sensory information*. Each sensory modality exhibits this distinction; pain circuitry is shown here. A) A nociceptive stimulus, such as heat above 45°C or cold below 25°C or sharp pricking pain, is detected by bare nerve endings and transported via Type A*δ* fibers toward the central nervous system. Dorsal root ganglion neurons (blue) impinge on spinal interneurons (grey). A polysynaptic pathway achieves a fast withdrawal reflex by exciting motor neurons which innervate flexor muscles in the ipsilateral limb and inhibiting motor neurons which innervate extensor muscles in the ipsilateral limb (orange); a crossed-extension reflex steadies the contralateral limb (not shown). Dorsal root ganglion neurons also send ascending fibers toward brain regions, including the periaqueductal grey (not shown) and the thalamus (grey). Thalamic neurons project to somatosensory cortical neurons (violet), where incoming signals are integrated with other inputs. B) Cortical neurons maintain high sensitivity to noise in gating the probability of firing a signal. An action potential is shown in green; once this event starts, it goes to completion. The refractory period and resting potential are shown in yellow; it is unlikely for a neuron to reach action potential threshold from this voltage, given only small noisy currents. A cortical up-state is shown in red; during this phase, noisy events contribute to signaling outcomes. C) Peripheral sensory neurons are not sensitive to random electrical noise in gating signaling outcomes; stimuli will produce a deterministic, stereotyped outcome. D) Peripheral motor neurons are not sensitive to random electrical noise in gating signaling outcomes; stimuli will produce a deterministic, stereotyped outcome.

Although both cortical neural circuits and spinal reflex circuits transduce sensory input into contextually-appropriate motor output, only cortical neural circuits generate qualitative perceptual experience and nonreflexive behavioral outcomes. For example, damage to the cerebral cortex ablates perceptual content and stimulus-evoked behavior [22]. The conscious experience of pain is correlated with neural activity in cortical circuits [23] but not spinal reflex circuits [24]. This model asserts that a unique physiological property of cortical neurons is correlated with the generation of representative information content. The assumption here is that: If a network of computational units computes signaling outcomes deterministically, merely as the sum of all input signals over some spatiotemporal window, then it will not produce representational information content. If instead signaling outcomes for each computational unit are probabilistic, then those neural circuits will produce representational information content. It is this proposed key property – retaining sensitivity to probabilistic events while gating a state change in the computational unit – that is necessary for the neural circuitry to generate perceivable information content.

This property - of retaining sensitivity to random electrical noise in gating signaling outcomes - is unique to cortical neurons. Neurons in the invertebrate nervous system exhibit near-perfect reliability and efficiency in information transmission [13, 25]. Similarly, neurons in mammalian spinal reflex circuits exhibit highly efficient firing patterns which can be predicted simply by summing upstream inputs [12]. Yet these predictable signaling outcomes are not characteristic of cortical neurons [14, 15]. Stochastic events and spontaneous subthreshold fluctuations in membrane potential affect the likelihood of a cortical neuron firing [16, 17]. Permitting random electrical noise to contribute to signaling outcomes boosts inefficiency and error rate within digital communication channels, so any well-optimized binary computing system should be highly robust to such noise. But rather than positing that cortical neurons have evolved to be more sloppy and inefficient binary computing units than spinal reflex circuits, the model asserts that random electrical noise is being productively utilized in this context to make non-deterministic computations. As a result, cortical neurons are predicted to be more efficient and more reliable at processing information than peripheral neurons, despite retaining a sensitivity to noise.

Indeed, cortical neurons actively maintain ‘up-states’ – with cells across the network suspended near action potential threshold and permitting stochastic charge flux to gate a state change [26]. These coordinated up-states facilitate the synchronous bursting activity that is associated with wakeful awareness and sensory perception [27–29]. Here, it is specifically predicted that cortical neurons in up-state, residing near the threshold for firing an action potential, are harnessing random noise to drive a probabilistic computation. Meanwhile peripheral neurons, with more deterministic outcomes, are not expected to physically generate a probability distribution, or a physical quantity of information.

The physical and mathematical definition of information is the set of all possible system macrostates, or the mixed sum of all component microstates [30]. In any particle system, this complex-valued probability density accounts for all possible spatial positions and atomic orbitals of every component electron in the system, over some time evolution. Randomness or disorder increase this physical quantity of information, while consistencies or patterns decrease this physical quantity of information. In classical models, the neuron is a binary computational unit, like a transistor, always in an off-state or on state. In this new model, the cortical neuron physically encodes information, with the membrane potential being the mixed sum of all component electron states. As the probabilistic component states interfere, possible states are reduced and information is physically compressed. The process of integrating upstream signals with random electrical noise should generate a cohesive quantity of information, or von Neumann entropy, which comprises both incoming sensory data from the local environment and the constraints of the encoding system.

### E. Justification for modeling the neuronal membrane as a holographic recording surface

A second assumption is that the outer membrane of the computational unit is a charge-detecting surface comprised of organic or synthetic polymers, which meets the criteria for a holographic recording surface. The justification for this assumption is provided in Figure 2.

**Figure 2.**
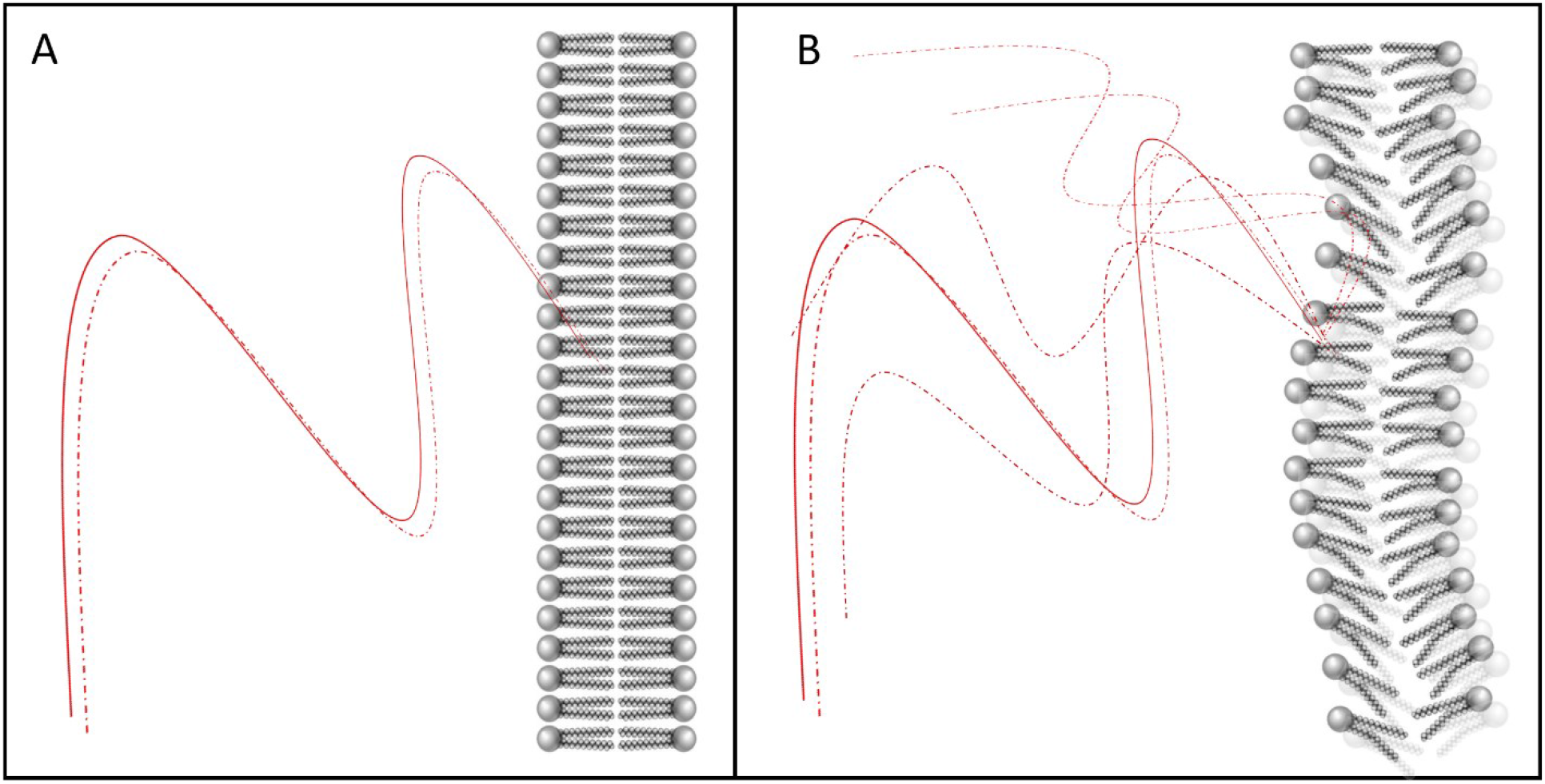
The unique anatomical feature of cortical neurons: a charge-detecting polymer surface. The proposed model asserts the outer plasma membranes of cortical neurons meet the criteria for a holographic recording surface: organic polymer surfaces with sufficient dimensionality to capture complex waves, a flat spatial frequency response, a large dynamic range, sensitivity to low energy exposures, high efficiency in coding, and protection from environmental factors. The electrical and mechanical properties of the neural membrane are dependent on lipid composition. (A) Increased fatty acid tail saturation is expected to increase lipid packing and the rigidity of the polymer surface. (B) By contrast, higher unsaturated fatty acid content is expected to increase the rate of membrane undulation. Neuronal membranes are comprised of different lipid species, compared with other cell types, and undulate over millisecond timescales more than other cell types [36]. The phospholipid composition of the membrane may also affect the electrical dipole moments induced by nearby electron-electron interactions (A, B), with phosphoglyceride content affecting dipole-induced membrane potentials. Notably, neural membranes contain different patterns of docosohexanoic acid-containing phosphoglycerides than other cell membranes, with these small structural changes leading to significantly higher membrane potentials [72]. The enormous diversity of lipid membrane composition (influenced not only by genetic and epigenetic factors, but also by dietary and lifestyle factors) are expected to influence the electrical and mechanical properties of the putative holographic recording surface, and therefore the qualitative properties of the reconstructed information content.

The physical process of holographic projection is well-established. Briefly, the position of an (n+1)-dimensional electromagnetic point source is physically encoded by a process of wave interference on an (n)-dimensional recording surface, generating a holographic projection of the encoded information content across (n+1) dimensions. Classical holograms generally encode information about each contributing electromagnetic point source on a two-dimensional surface, then project the rich, qualitative information content into a three-dimensional volume [31, 32]. This method can be extrapolated to describe any electromagnetic point source impinging on any holographic recording surface [33, 34].

The criteria for an ideal holographic recording surface are: 1) a charge-detecting surface comprised of organic or synthetic polymers, with 2) a linear relationship between energy exposure and the amplitude of the reconstructed wave, attaining signal fidelity; 3) a flat spatial frequency response, ensuring signal capture; 4) a large dynamic range, providing a good signal-to-noise ratio; 5) a high-quality and lossless material, affording efficiency in projecting the hologram; 6) sensitivity to low energy exposure, achieving fine signal detection; and 7) protection from environmental factors that impact functionality.

These criteria are met in the case of a biological neural network that retains sensitivity in detecting random electrical noise. Every nearby electron is an electromagnetic point source, with an inherently uncertain spatial position and atomic orbital; these complex-valued probability amplitudes undergo wave-like interference. The putative encoding surface is a gelatinous phospholipid bilayer comprised of organic polymers, whose shape is described along three orthogonal spatial axes [35]; whose shape is also changing over time [36]; and whose electrical properties permit high-fidelity signal capture over a large dynamic range.

Any information encoded on this surface by the physical interference of wave-like particles must be projected into a higher dimensionality, upon interference with a reference beam, in accordance with the holographic principle. This model yields a multi-sensory percept, representing the spatial location and intensity of all attended stimuli within the receptive field at that moment.

### F. Modeling each electron as a complex wave

Classical approaches in neuroscience describe each neuron as being in either ‘on-state’ or ‘off-state’ at any given moment, like a transistor. By contrast, this new approach describes every cortical neuron nearing action potential threshold as having some probability of switching from an ‘off-state’ to an ‘on-state’. The probability of a cortical neuron reaching action potential threshold is dependent on the spatial position and energy state of every component electron. The state of each electron remains uncertain in the context of a dynamically-changing electrical field, which is affected by every neural membrane in the vicinity. And so, in this model, each neuron acts as a qubit, rather than a classical binary computing unit, encoding the probability of firing an action potential as a function of all component electron states. Any neuron in an uncertain state, with some probability of either firing or not firing, will contribute to the probabilistic neural network state, while neurons with purely deterministic signaling outcomes will *not* contribute to that density of possible system states. The mixed sum of all component states is the amount of information held by the system, so the mixed sum of all contributing microstates yields the quantity of information encoded by a given neuron and the mixed sum of all contributing neuronal states yields the quantity of information encoded by the network.

Since the properties of electrons are undefined in the present moment (including how much time has passed to reach the present moment), the spatial position, temporal location, and atomic orbital of an electron are best described as complex-valued probability amplitudes (Figure 3). After some time evolution, the spatial position of each electron is a scalar multiple of the Planck length, with respect to its previous position; the amount of time passed is a scalar multiple of the Planck time; and the energy state of each electron is a discrete orbital configuration. Since each electron within the system exists in a probabilistic state, with an uncertain trajectory, each electron is best described as a complex-valued probability amplitude defined along five orthogonal axes – namely *x, y, z* (spatial position), *t* (the amount of time passed), and *ω* (the amount of energy above ground state, which is expended to create information). The total energy expended by the system toward creating the probability density is the Hamiltonian: the total amount of energy available for redistribution across the system. Constructive and destructive interference of these complex-valued probability amplitudes yields a non-deterministic resolution of the present neural network state, such that each electron is assigned a discrete state. The voltage state of each neuron is resolved instantly, as every component pure state is resolved. The computational cycle then begins again, with a new probability density emerging. Any energy distributed toward entropy over the lifetime of a non-dissipative thermodynamic system, which is not recovered during a computation, remains distributed along the ‘time’ and ‘energy’ axes.

**Figure 3.**
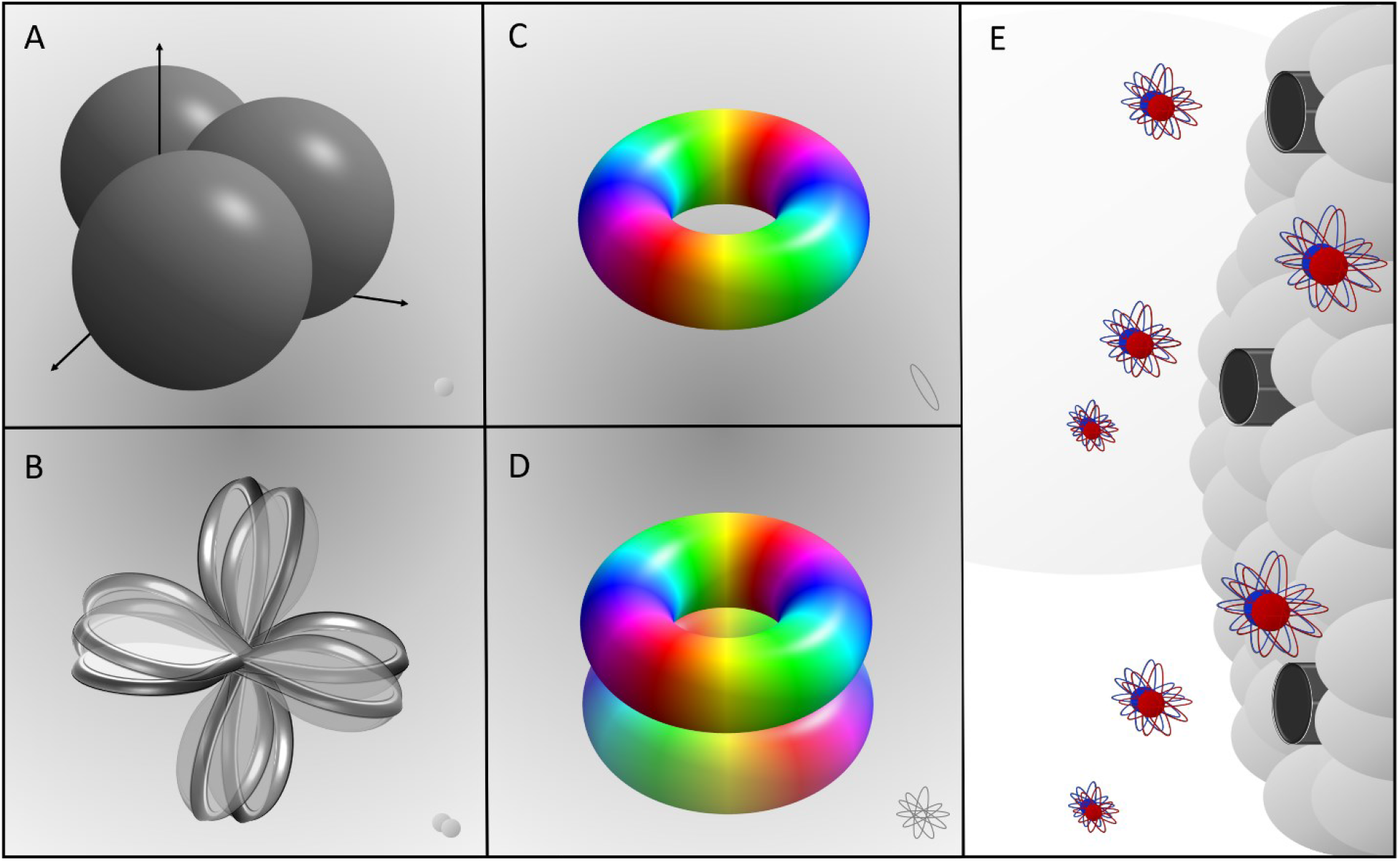
A schematic of probabilistic electron interactions with the neural membrane. A) Each electron has some position across the *x, y*, and *z* axis, relative to the neuronal membrane surface. Each of these position values is uncertain, described by probability density rather than a defined state. As a result of the forces between electrons and the nucleus, and interactions of each ion with other atoms in the vicinity, an entire ion also has some probabilistic position (right corner). B) The *x*, *y*, *z* position of an electron changes over time *t*, as the system is perturbed by its environment. C) The energy state of an electron *ω* is also uncertain, best described as a probability density rather than a defined state. D) The likely energy state of an electron also changes over time *t*, as the system is perturbed by its environment. E) Each electron, acting as an electromagnetic point source, impinges probabilistically on each nearby neuronal membrane surface. The previous state of each electron, at *t*_0_, serves as a reference beam (red); any change in *x*, *y*, *z* position or energy orbital ω over time *t* yields an object beam (blue). Each object beam, representing a possible electron state, interferes with the reference beam, thereby encoding information of the surface of the neuronal membrane. Given the uncertainty of each electron state, the neuronal membrane potential is uncertain in the present moment (unless the cell is in a refractory state or has already reached action potential threshold). In this model, the trajectory of each electron is a complex-valued probability amplitude described across five orthogonal axes (*x*, *y*, *z, ω* and *t*); the interference of these complex-valued probability amplitudes on the polymer neural membrane surface physically encodes information.

### G. Electromagnetic point sources can be modeled as interfering object and reference waves on a two-dimensional holographic recording surface

Classical holograms are formed by the interference of light waves, with the information encoded on a two-dimensional surface and projected into a three-dimensional volume [32, 33]. It is worth noting that these principles can be extrapolated to any wave-like particle, to generate a hologram of the electrical field [44]. A photon, an electron, or even an entire ion, can be represented as the point source of an electromagnetic field, with the amplitude and phase of the wave represented by its absolute value and relative angle. Once the wave is split into a reference beam (which undergoes no change) and an object beam (which undergoes some change in position over some period of time), the intersection of the two beams causes a phase shift, given by *ϕ*:

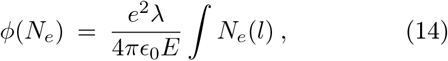

Where:

*N_e_* is the density of electrons or light waves, integrated over a path of length *l* over time *t*;
λ is the wavelength of the electromagnetic point source; e is the charge of the electromagnetic point source;
*E* is the energy of the electromagnetic point source; and *ε*_0_ is the permittivity of free space.

The reference beam and the object beam will interfere on the holographic recording surface. Alone, the reference beam and the object beam are expressed in terms of their respective magnitude and phase [45], given by:

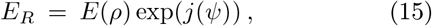

And:

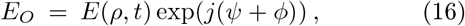

Where:

*E_R_* and *E_O_* are the optical or electrical fields of the reference and object beams, respectively;
*ρ* is the position of the electromagnetic point source; *ψ* is a constant defined by 2*π*/λ;
*ϕ* is the effective phase angle between the object and reference waves after elapsed time *t*;
and *j* is the imaginary part of the complex wave.

The spatial distribution of a point source with strength *E* can then be modeled as a spherical wave [46], given by:

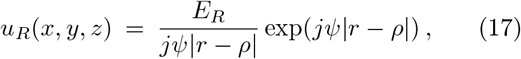

And:

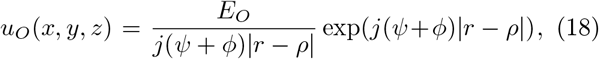

Where:

*ρ* is the position of point *E*, given by 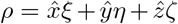;
*r* provides the coordinates of the recording plane, given by 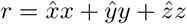;
the ‘hat’ denotes any unit vector along the identified coordinate plane;
and *ξ*, *η*, and *ζ* are coordinate values along the *x, y*, and *z* axes.

The origin of the encoded signal is an electromagnetic point source, generating a wave propagating toward the recording surface along a single axis. The distribution of this wave on the holographic recording surface is the difference between *r* and *ρ* [46], given by:

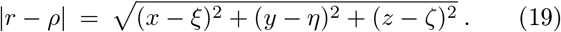

However, if the recording plane is a two-dimensional surface, then *z* = 0 and the equation simplifies:

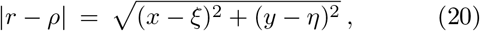

With the paraxial approximation given by:

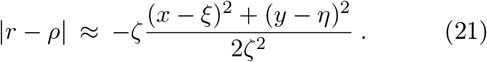

As the beams intersect on a two-dimensional holographic recording surface, they encode the object in a pattern of wave interference. The intensity *I* of the combined beams is proportional to the square of the combined waves or the square of the magnitude of the electrical field:

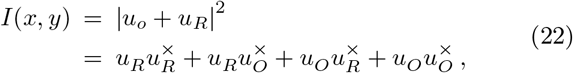

Where:

× indicates the complex conjugate of the written value. Here, the first and fourth term constitute zeroth-order diffraction and do not contribute to the reconstructed image. Meanwhile, the second term encodes the true image, and the third term encodes a phase-conjugate or virtual image, which is spatially separated from the real image represented by the second term. Together, the virtual and real images will combine to form a hologram with increased dimensionality compared to the holographic recording surface that encodes it.

The overall transmittance *T* is related to the sum of all intensities *I*. This value is found by integrating every combined beam impinging on the holographic plate, and adjusting for key factors related to the properties of the holographic recording plate:

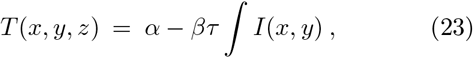

Where: *α* is a constant which accounts for the uniform background electrical field;

*β* is a constant measuring the sensitivity of the surface to an electrical pulse of wavelength λ;

and *τ* is the duration of exposure of the surface to the electrical pulse of wavelength λ.

### H. Information encoded by interfering waves on a two-dimensional recording surface can be reconstructed into a three-dimensional volume

Combining the transmittance with the reference beam yields a reconstructed electrical field *E_rec_*, describing the spatial distribution of the now-diffracted electrical field strength:

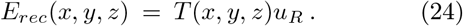

Substituting Eqs. 22 and 23 into Eq. 24 yields the full equation. Here, the reconstructed electrical field is described in a higher-dimensional coordinate plane, as a function of the reference beam interacting with the recording plate. This causes the diffraction pattern to be holographically projected, with a resolution determined by the properties of the recording surface:

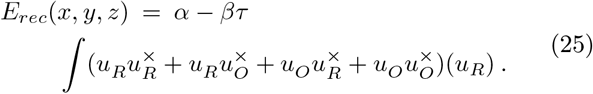

The interaction of the reference beam with the twodimensional holographic recording surface generates a three-dimensional image, containing the zeroth-order diffraction (essentially the original reference wave) and first-order diffraction (including the object beam, multiplied by a scalar, and the object beam conjugate, which provides the reverse curvature of the object beam). It should be noted that these laws can be extrapolated into any dimensionality, with the mathematics of wave interference and the resulting diffraction pattern merely increasing by an order of complexity. The only requirement is a holographic recording surface of sufficient dimensionality to encode the charged particles or electromagnetic point sources as complex waves.

## III. RESULTS

### A. Electrical noise and upstream signals are encoded on a three-dimensional recording surface

Each electron has some probabilistic spatial location and energy state, relative to each neuronal membrane. Electrons outside the brain have very little probability of affecting the membrane potential of any neuron, so only electrons within the system are included when modeling the membrane potential as the mixed sum of all component pure states, or the information entropy held by the network. Electrons do not have defined states in the present moment, but rather have some range of possible positions *r*, momenta *s*, and energy states *ω* [47, 48]. Each electron is described by a wavefunction that provides its likely positions during this time evolution, generating both a reference path (with no interactions) and any number of object paths (with interactions). Each electron can therefore be modeled as an object with a probabilistic position defined across *x, y*, and *z* axes, with an atomic orbital *ω* at some time *t*. All five of these values are intrinsically uncertain; *x*, *y*, *z*, *ω*, and *t* are scalar multiples of Planck units mapped along five orthogonal axes (Figure 3). The probabilistic state of each electron is therefore best described as some set of possible trajectories, relative to each neuronal membrane surface.

Each electron is described by a reference beam *E_R_* and one or more object beams *E_O_*. *E_R_* is separated from each possible value of *E_O_* by some phase angle, denoted by *ϕ*, which is generated by the uncertain state of each electron. The electrochemical potential of each neuronal membrane is dependent on the position of each electron in the vicinity, and that voltage state is a probabilistic function of all possible component pure states. The state of each computational unit is resolved as each component microstate is resolved, with the reference beam *E_R_* of each electron interfering with every possible object beam *E_O_*. The equations describing *E_O_* and *E_R_* are identical to Eqs. 15 and 16 given above. The spatial distribution of a point source with strength *E*, impinging on a threedimensional recording plane, can then be modeled as a spherical wave, given by:

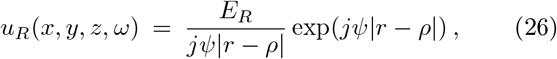

And:

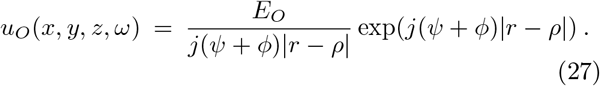

The relationship between the *x, y, z* location of each electromagnetic point source and the coordinates of the holographic recording surface, after time *t*, is given by:

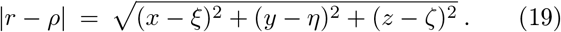

But this value cannot be reduced or approximated as before. Because the neural membrane is a charge-detecting polymer surface structured across a threedimensional coordinate plane, *the entire neural network acts as a three-dimensional holographic recording plate*, encoding the interfering reference beams and object beams of many nearby electrons simultaneously. Any charge crossing this surface will be detected as a shift in the electrochemical potential of the neural membrane.

To describe the interference pattern for a given electron impinging on a biological neural network structure, the *z* axis is not negligible and must be included in the calculations; the interference patterns of *E* on the holographic recording surface must include coordinates across the *x, y*, and *z* dimensions. As the reference and object beams interact with a three-dimensional neural membrane, a hologram is encoded in a pattern of wave interference between the reference beam and all possible object beams for each electron. The intensity *I* of the combined beams is proportional to the square of the combined waves or the square of the magnitude of the electrical field:

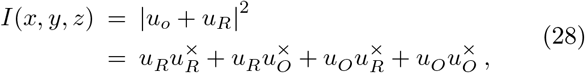

But now, the intensity of the combined beams on the recording surface cannot be simplified to two dimensions; it is defined along three axes. The transmittance *T* achieved by the three-dimensional holographic recording surface is related to the intensity *I* for each combined beam impinging on that surface. Integrating all of these intensities yields a transmittance of the four-dimensional spherical waves:

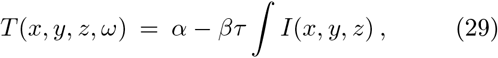

Combining the transmittance with the reference beam yields a reconstructed electrical field *E_rec_*:

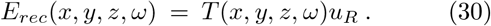

Substituting equations once again permits the reconstructed field to be described in a higher-dimensional co-ordinate plane. This volume is a function of the sum of all possible trajectories, with regard to a reference beam and the diffraction pattern on the holographic plate:

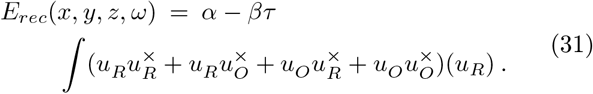

The result, as before, is to generate a holographic reconstruction in a higher dimension than the holographic recording surface. In the case of biological neural networks, the interference occurs between a reference wave (denoting non-movement of a given electron over some period of time) and an object wave (denoting movement of that same electron over that same period of time). This interference is encoded on the three-dimensional neural membrane structure, as a charge is detected. Each point source is represented by both the reference wave and every possible object wave. The resulting interference generates a combined holographic representation of the trajectory of every electron in the system. During this process of interference, the voltage state of each neuron is defined as every component pure state is defined. Any data encoded by that system macrostate is naturally paired with projected information content, exclusively accessed by the encoding structure, as long as that encoding structure meets the necessary criteria for a holographic recording surface.

### B. Non-deterministic computational outcomes emerge from probabilistic interference

The encoding process generates a holographic reconstruction of information in its original dimensionality. The interference pattern that results from this physical computation is predicted to amplify high-likelihood neural network states while canceling out noisy states that are not compatible with the reference wavefunction. Thus, multiple object waves interacting with a reference wave may lead to a new ‘confidence’ in the most probable present state for the entire system. This can also be understood as a contextually-relevant mental model ‘informing’ the most probable present state for the entire system, by acting as a reference beam or reference state. This compression event can be calculated by taking the derivative of all possible values for the phase angle *ϕ*, with respect to all factors that contribute to the distribution of probability amplitudes λ. Taking the derivative identifies the surface boundary of the high-dimensional volume of probability, thereby reducing a wavefunction density into a single observable system state:

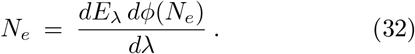

The integrated neural network state in the present moment is represented by a hyperspherical wave or wave-function. The reference wave, *u_R_*, is identical to the immediately previous neural network state. The object wave, *u_O_*, has experienced some change – and again, there are many possible object waves here. The interference pattern between the reference beam and all possible object beams can be considered as a function of the phase angle difference between the previous neural network state and the current neural network state. Equation 32 thus describes an information compression event, with the constructive and destructive interference between complex-valued probability amplitudes yielding a non-deterministic computational outcome. Every electron is assigned some position and energy state, before embarking on a new uncertain trajectory. In other words, taking the derivative of that volume of information content is equivalent to identifying observable values on the boundary region. It is worth noting that Equation 32 is equivalent to the Hellman-Feynman equation:

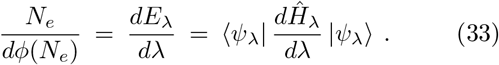

Here, the total energy of the system E relates to: the Hamiltonian operator *Ĥ*_λ_, any perturbing factor λ that contributes to the expanse of the probability distribution, and any eigenvalue *ψ*_λ_. Eigenvalues can only be observed at the beginning and the end of a computational cycle; they are otherwise undefined. Dividing the volume of probability by the derivative of the phase angle *φ* (which denotes a change in the system-wide state from the previous or reference state) is equivalent to dividing the derivative of the total energy of the system by the derivative of any factors contributing to the expanse of the probability distribution. In the Hellman-Feynman equation, this relationship is also equal to the path taken through the probability distribution to arrive at some observable *ψ*_λ_ [47].

Thus, any change from the reference neural network state must involve a reduction of the probability density, reducing the number of possible neural network states. The system undergoes a non-deterministic computational process to reduce the expanse of possible system states into the most likely and consistent state. In practical terms, the system state is restricted to compatible and permitted states for all participating particles within the system, which cannot occupy the same spatial position, spin, and energy state. The value of the derivative of the phase angle must provide a compatible and permitted state for all particles in the system. In the interim, the system produces a qualitative holographic reconstruction of the ‘information’ it has encoded. These wavefunctions not only generate perceived content, but also constructively and destructively interfere with the reference wavefunction to realize the most likely and consistent state for every particle in the system.

In short, the integration of probabilistic particle behavior across a three-dimensional neural network generates a combined wavefunction, describing all possible states for the system. This probability density is the mathematical definition of information. The encoded information is naturally projected into a higher-dimensional volume, in accordance with the holographic principle. The timescale of this coherent state should provide a ‘frame rate’ at which perceptual experience is updated.

The holographic recording surface, which is described in three spatial dimensions, is *itself* also changing over time, as neuronal connections grow into a more ordered state [49]. This remodeling process changes the mass and composition of each synapse. That is: the mass distribution of the system also changes over time. Individual neural network states - each paired with qualitative information content - are integrated into temporal sequences, by integrating information (or system macrostates) along the *mass* axis. This integration of a sequence of neural network states provides a high-dimensional map of how *both mass and energy* have been distributed over the lifetime of the non-dissipative thermodynamic system. Temporal sequences of neural network states should be paired with temporal sequences of perceived events, which predict or represent the cause-effect structure of reality.

### C. Specific predictions of this model

This approach usefully provides a mechanism by which cortical neural activity generates perceivable information content. Since cortical neurons allow random electrical noise to gate a signaling outcome, and each electron may contribute to the state of multiple neurons, the state of each neuron must be computed simultaneously, as the state of each component pure state is resolved. If this system-wide non-deterministic computation involves physically encoding information, in the form of complex waves, on a polymer recording surface, the information will be perceivable to the encoding system. A number of specific predictions arise from this model, providing the opportunity to empirically test the proposed mechanistic relationship between cortical neuron anatomy and physiology with qualitative perceptual experience.

#### 1. The predicted neurophysiological correlates of perception

In this new theoretical framework, each neuron residing in an uncertain state contributes to a system-wide probability distribution, or the quantity of physical information held by the encoding system. So, while all particle systems are characterized by probabilistic particle behavior, only macroscale systems which encode these probabilistic events into the voltage state of a macro-scale computational unit will capture that probabilistic particle behavior. Any biological or engineered system which operates on these principles, characterized by cortical up-states in mammals, pallial up-states in birds, or analogous up-states in (hardware-instantiated) engineered neural networks, should exhibit a capacity for spontaneous, unprogrammed, non-deterministic but contextually-appropriate behavioral output. Purely deterministic circuits, which do not allow random electrical noise to affect signaling outcomes, may be able to process sensory inputs and motor outputs, but should only achieve stimulus-driven reflexive action.

Since the encoding of information is predicted to be correlated with information content, trajectories of electromagnetic point charges impinging on the neural membrane should define the content of the perceptual experience. The unique qualitative nature of the holographic projection should therefore rely on several factors: the recent trajectory of each ion, which contributes to the signaling outcome of each neuron; the ion conductances and opening kinetics of each channel; neuronal morphology and local membrane permeability; the electrochemical potential of the neural membrane, which is dependent on these factors; and finally, the physical location of the organism, which makes certain observations possible, and the organism’s ability to notice these sensory stimuli, given their contextual expectations and the amount of energy they are devoting to attending a stimulus. This model predicts that encoding probabilistic charge flux into the voltage state of a computational unit provides the physical basis for perceptual experience. Experimentally manipulating any of the above factors should alter the perceptual content encoded by that neuronal population and should affect contextually-dependent behavior.

#### 2. The predicted neuroanatomical correlates of perception

In this new theoretical framework, each neuron in an uncertain state contributes to a system-wide probability distribution, or the quantity of physical information held by the encoding system. So, while all cells in the body have polymer membranes, only those capturing probabilistic particle behavior as interfering complex waves may encode and project information. And only if that charge-detecting polymer membrane meets the criteria for a holographic recording surface, will the information content be perceivable to the system encoding it; the holographic projection of the complex wavefunction is hypothesized to be equivalent to perceptual experience. Any biological or engineered system which operates on these principles, characterized by cortical up-states in mammals, pallial up-states in birds, or analogous upstates in engineered neural networks, should be capable of having perceptual experience. Again, by contrast, purely deterministic neural circuits should achieve only stimulus-driven reflexive action, without any corresponding perceptual content.

Since the physical encoding of information is predicted to be correlated with information content, the properties of the neural membrane should also define the content of the perceptual experience. The unique qualitative nature of the holographic projection should therefore rely on several factors: the quantity and distribution of cholesterol molecules within the neural membrane, and the complement of polyunsaturated fatty acids comprising the neural membrane, which both affect both the rigidity and the electrical properties of that putative holographic recording surface. Experimentally manipulating any of the above factors should alter the perceptual content encoded by that neuronal population and should affect stimulus-dependent behavioral choice in perceptual tasks.

#### 3. The predicted effects of electromagnetic stimulation

Direct electrical stimulation of cortical neurons should affect firing rates, but exogenous electrical fields or the mere presence of metal objects should have no effect on cortical neuron activity or perceptual content. The probability density associated with the physical state of the neural network should not change, unless the electrical pulse affects the probabilistic state of some neurons but not others. Therefore, this theory predicts that electrical stimulation directed at a population of cells should prompt neuronal firing and alterations in perception, but exogenous electrical fields and metal objects should have no discernable effect.

Magnetic stimulation should exert an effect on neuronal activity, if directed toward a subset of cells within the cortex, but uniform magnetic fields encompassing the entire brain should have no effect. The latter experimental manipulation will not affect the density of possible system states but rather warp all particle trajectories equally. The information content encoded by the neural network should not be altered, unless the magnetic stimulation specifically affects the probability of neuronal firing. Therefore, this theory predicts that magnetic stimulation directed at a region of cortex should prompt changes in neuronal activity and perceptual content, but exogenous magnetic fields such as those exerted by head-surrounding magnetic resonance imaging equipment should have no discernable effect.

#### 4. The predicted effects of specific drug classes

General anesthetics, alcohol, barbiturates and benzodiazepines enhance GABA signaling and reduce perceptual awareness [50–52]. GABA receptor agonism leads to sustained inhibitory potentials in cortical neurons, significantly decreasing the likelihood of a signaling outcome [53, 54]. This leads to a significant reduction in gamma frequency oscillations across neocortex [55]. This theoretical framework agrees with classical neuroscience on the mechanisms underpinning neural computation, but goes a step further than classical neuroscience to describe how reduced neural activity effectively leads to motor slackness and decreases in perceptual content. Here, enhanced GABA signaling reduces the density of possible system macrostates, and therefore reduces the holographic projection of information content. This theory predicts that increasing excitatory post-synaptic currents should reverse the effects of GABA receptor agonism, restoring both motor activity and perceptual content.

Glutamate receptor agonism drives cortical neuron activation, converting neuronal signaling from a probabilistic outcome to a more deterministic outcome; but conversely to GABA agonism, cortical neurons are forced to fire an action potential rather than rendered less likely to do so [56]. So, enhancement of glutamatergic signaling with application of NMDA or AMPA receptor agonists should lead to rigid high-frequency patterns of neural activity and a corresponding reduction in perceptual awareness. Indeed, any experimental manipulation that makes neural signaling outcomes more deterministic across neocortex should be correlated with decreased perceptual content and motor rigidity. This theoretical framework agrees with classical neuroscience on the mechanisms of neural computation, but goes a step further to describe how excessive glutamatergic signaling leads to motor rigidity and a decrease in the richness of perceptual content. Here, a reduction in the density of possible neural network states reduces the holographic projection of information content and the range of possible behavioral outcomes resulting from cortical computation. This theoretical model therefore predicts that boosting inhibitory post-synaptic currents should increase the probabilistic nature of cortical neuron activity and should reverse the effects of glutamatergic over-excitation, restoring both perceptual content and motor control.

Hallucinogens such as LSD alter perceptual experience [57]. In classical neuroscience, there is little understanding of how this class of drugs leads to changes in mental representation. Here, an increased uncertainty in signaling outcomes, rather than any particular pattern of activity across neuronal populations, is expected to underlie the enhanced quality, quantity, and binding of perceptual experience. If uncertainty can be sustained for longer timescales, the expanse of possible neural network states is increased; in such cases, more information content should be generated, and more perceptual experience should be available to the neural network. This theory predicts that drugs like LSD or psilocybin, which enhance perceptual experience, do so by rendering neural activity more probabilistic. As a result, it is expected that many more EPSCs and IPSCs will occur during upstate, prior to cortical neurons achieving threshold for firing an action potential, in this pharmacological context. This class of drugs is therefore predicted to be therapeutically useful: Increasing the probabilistic distribution of neural network states permits the selection of alternatives. This process may help some individuals to overcome unhealthy behaviors that rely on habitually-ingrained patterns of neural activity, such as addictions, by thermodynamically favoring alternative patterns of neural activity and the adoption of alternative routines in behavior.

## IV. DISCUSSION

The holographic principle is a key property in information theory and modern physics [58]. This principle states that information encoded on a lower-dimensional boundary region is paired with a higher-dimensional volume of information content. That volume is defined by the Bekenstein bound, which provides that the maximum density of information is proportional to the surface area of the system [59]. The present report systematically explores how natural computational systems may obey the holographic principle, by both physically encoding and representing information. The criteria required to achieve non-deterministic computation and to generate perceivable information content in this model are listed in Table 1.

**Table 1.** Listed are the criteria for a polymer substrate to act as a holographic recording surface and the criteria for a macro-scale system to achieve quantum computation. The computational unit must have a probabilistic state, due to coulomb scattering. This uncertainty must be productively sustained until component states resolve, yielding observables or eigenvalues. Random fluctuations should observably contribute to signaling outcomes. If all component pure states within the neural network are interdependent, any one outcome will exclude other possible outcomes. The physical quantity of information is encoded by the anatomical and physiological properties of the computational unit. The information content encoded will only be perceivable by the system if the computational units meet the specific criteria for a holographic recording surface (* and have sufficient dimensionality to capture high-dimensional wavefunctions). If all criteria are met, the system will produce a cohesive holographic projection of information content during the non-deterministic computation.

| Criteria for an effective quantum computing system |  |
| --- | --- |
| Characteristic | Function |
| broad coulomb scattering of ions on the surface of the computational unit | sustains uncertainty |
| coherence timescales of ions are longer than thermal dissipation timescales | sustains uncertainty |
| the state of each ion is dependent on the state of other ions in the system | creates cohesive information |
| signaling outcomes are dependent on both upstream inputs and noise | achieves probabilistic computation |
| Criteria for an effective holographic recording surface |  |
| Characteristic | Function |
| a charge-detecting surface comprised of organic or synthetic polymers | encodes wave interference |
| a three-dimensional surface capable of structural self-remodeling over time | encodes particle interference * |
| signals are faithfully transduced into the amplitude of a reconstructed wave | retains signal fidelity |
| surface has a flat spatial frequency response | ensures signal capture |
| surface has a large dynamic range | improves signal-to-noise ratio |
| surface is a high-quality and lossless material | affords highly efficient coding |
| surface is sensitive to low energy exposure | attains fine signal detection |
| surface is protected from environmental factors that could affect its integrity | preserves continuous operation |

This new approach focuses on the unique properties of cortical neurons. For neurons to physically generate and compress information, using this computational process to drive signaling outcomes, probabilistic particle behavior must be sustained for sufficiently long time periods to contribute to thermal fluctuation-dissipation dynamics at the neural membrane [37]. Furthermore, the system must be able to recover some of the free energy initially expended on information generation during information compression [38–41], and must be able to use this newly-released thermal free energy to drive macro-scale outcomes [42]. Certainly, a number of researchers have expressed doubt that biological neural networks could be quantum systems, largely on the hypothetical grounds that quantum effects would be difficult to sustain in organic systems at the macro-scale [43]. This critical view is not supported by evidence, since neurophysiological studies readily demonstrate that random electrical noise contributes to cortical neuron signaling outcomes [16, 17].

This model employs two assumptions, which are empirically valid: 1) cortical neurons permit probabilistic events to affect the likelihood of firing an action potential, and 2) the membrane of each computational unit is a charge-detecting polymer substrate that meets the criteria of a holographic recording surface. If both criteria are met, then the encoding process will generate a holographic projection of representative information content. Since the probabilistic trajectory of each electron can affect the voltage state of multiple computational units, the system state must be computed as a whole, with the macrostate of the network defined as the state of each computational unit is defined and the macrostate of each computational unit defined as every component microstate is defined. The mixed sum of all component microstates is the information physically held by the encoding system. That information content is exclusively accessed by the system that generated it and is limited by the range and sensitivity of each contributing sensory apparatus. This model therefore offers the first plausible explanation for cortical neural activity being paired with qualitative perceptual experience.

A number of theorists over the past century have pointed to the unitary and experiential nature of mental representation [60, 61]. In the early twentieth century, the singular and irreducible nature of psychological experience was articulated by gestalt theorists Wolfgang Köhler and Maurice Merleau-Ponty, who differentiated between external reality, the act of sensation, and the experiential nature of perception [62, 63]. More recently, the concept of binding information within the time domain has become critical to studying the neural correlates of perception, memory, decision-making, and context-dependent cognitive expectations [64]. Indeed, these features of conscious awareness are associated with synchronous activity across sparsely-distributed neural populations [65]. Gamma frequency oscillations in particular have been proposed to be correlates of perceptual awareness [66]. However, the exact physical mechanisms by which neural signaling might generate temporally-bound qualitative information content have been difficult to articulate.

Holography provides a well-understood method by which discrete packets of information are bound into a cohesive whole, and this relevant prospect has been appreciated for some time. The inventor of holography Dennis Gabor, the quantum physicist David Bohm, the neuroscientist Karl Pribram, and the cognitive scientist Christopher Longuet-Higgins considered holography a potentially useful metaphorical framework for describing the unitary nature of consciousness. They pointed out that memory is a non-localizable phenomenon [67] and that Fourier transforms of electrical oscillations reproduce natural inter-spike intervals [68, 69]. However, they did not specify any mechanism or process by which neural networks could physically produce holographic information content, and so this view has remained a vague concept rather than a formal scientific theory. These mid-century gestalt theories, with no clear mechanistic basis, have been largely replaced by the framework of parallel distributed processing, which explains how complex pattern detection and decisional outputs can emerge naturally from neuronal ensembles [70, 71]. This minimalist view, standard in both cognitive sciences and computer engineering, remains the gold standard for the parsimonious accounting of neural computation. And yet, this approach does not explain the qualitative, experiential nature of perception; it also requires that neural networks are not structured in a completely random manner, but rather start out with some initial connectivity suited to the task at hand. A better model, one that captures the emergent properties of biological systems, is needed.

Here, modeling the probabilistic behavior of electrons at the neural membrane generates a holographic projection of information content. The hologram is encoded by the distribution of electromagnetic point sources impinging on the surface of neurons. The mixed sum of this probabilistic particle behavior over some period of time t encodes the likelihood of a state change in the neuron. The distribution of trajectories yields the set of possible system states; this physical quantity of information is encoded in the membrane potential of each neuron, such that each computational unit has some probabilitity of reaching action potential and firing. In this model, a neural network encodes information by sustaining a distribution of probabilistic states.

The neural network then computes the most internally-consistent system state from that probability density, by allowing the constructive and destructive interference of complex-valued probability amplitudes to identify a mutually compatible state for all component electrons. The transiently-defined system state, at some location in time, is paired with a holographic projection of representative information content. Each of these ‘snapshots’ of the physical neural network state, paired with qualitative information content, has some address along the time axis. Integrating these neural network states into temporal sequences causes interference between complex-valued probabilities, allowing the most likely sequence of system states to emerge. Here, a single neural network state encodes a multi-sensory percept, or a predictive semantical statement about the present state of the world, while a sequence of neural network states, each with some location along the time axis, encodes a cause-effect model, or a predictive syntactical statement about the world.

## V. CONCLUSIONS

This report models the behavior of individual electrons as electromagnetic point sources impinging on the neural membrane. If the probabilistic trajectory of each electron can affect the voltage state of multiple computational units, then the system state must be computed as a whole, with the state of the neural network being defined as each component pure state is defined. If the outer membrane of each computational unit also meets the criteria of a holographic recording surface, then the encoding process must also generate a holographic projection of representative information content. If these two empirically-valid anatomical and physiological criteria are met, then a neural network should produce both representative information content and non-deterministic signaling outcomes. The representative information content should correspond to data collected from the local environment by all available sensory modalities, should be accessible *only* by the encoding system, and should be limited by the range and sensitivity of the sensory apparatus. In this model, these emergent features are natural by-products of cortical information processing.

This model rejects dualism and posits that consciousness is a physical process, obeying physical laws – a process of neural computation. If the predictions of this theory pan out, and altering the anatomical properties of the encoding structure does change the content of perceptual experience, it might be sensible to assert a causal relationship between probabilistic neural activity, information generation, and mental representation. If the compression of information is correlated with ‘comprehension’, and interrupting this physiological process interferes with synchronous neural activity and subsequent behavior, it might be sensible to assert a causal relationship between probabilistic neural activity, information compression, and motor output. A preponderance of evidence will be needed to discard the null hypothesis, and confidently assert that neural activity does generate a holographic projection of information content, with the interference of complex-valued probability amplitudes driving a non-deterministic computation.

## ACKNOWLEDGMENTS

The author received support for this work from the Western Institute for Advanced Study, with generous donations from Jason Palmer, Bala Parthasarathy, and Vanguard Charitable.

3D rendered images were produced by the python script hydrogen, which was written by User Geek3 and shared under a creative commons license.

## Notes

### Competing Interest Statement

The authors have declared no competing interest.

